# The clone wars: the Japanese knotweed (*Fallopia japonica*) vs the black vine weevil (*Otiorhynchus sulcatus*) – characterization of a potential herbivory

**DOI:** 10.1101/2022.09.27.509672

**Authors:** Loic Teulier, Sara Puijalon, Christelle Boisselet, Florence Piola

**Author notes:** **Correspondence:** Loic TEULIER and Florence Piola.

## Abstract

The Japanese knotweed (*Fallopia japonica*) is considered as highly invasive in Europe and is largely widespread in France, without any established predator. This short study first characterized the herbivory of *Fallopia* by the black vine weevil (*Otiorhynchus sulcatus*), a commonly encountered coleopteran in France. Through an experimental design of leaf choices, between *Fragaria* spp. and *Fallopia* spp., our results show that the insects prefer *Fallopia*, even if it is presented for the first time. Even if this simple observation may appear as trivial, it highlights a novel plant-insect interaction and may start new insight in plant control or invasion management.

## Introduction

Japanese knotweed [*Fallopia japonica* (Houtt.) Ronse Decraene var. *japonica*, Polygonaceae, hereafter *F. japonica*] is native to Japan and Eastern Asia (Barney et al. 2006). It was introduced into Europe in the 19th century by Phillipe von Siebold (Bailey and Conolly 2000) and is now widespread in North America and Europe (Bailey et al. 2009; Rouifed et al. 2014; Gippet et al. 2018). This species is a vigorous herbaceous perennial and to date, in Western Europe (i.e. Germany, Switzerland, United Kingdom, Belgium and France), only a single octoploid female (male-sterile) clone has been reported (Buhk and Thielsch 2015; Hollingsworth and Bailey 2000; Krebs et al. 2010; Tiébré et al. 2007).

In its country of origin, *Fallopia japonica* is eaten by some specific insects, such as the Japanese beetle, *Popillia japonica* (Kawano et al. 1999) or the Japanese Knotweed psyllid, *Aphalara itadori* (Shaw et al. 2009). As the former was identified as a crop pest in North America, the latter is currently experimented as a potential biocontrol agent in UK. Other candidates, such as the Common Amber snail, in Slovenia, tried to be identified as native biological agent to control the widespread of *Fallopia* spp. (Laznik and Trdan, 2017). And despite a high interest for controlling this invasive plant species (Cottet et al. 2015), to our knowledge, there is still a lack of identified enemy of the Japanese knotweed in France. (Maurel et al. 2013). However, we have fortuitously observed an important herbivory of *Fallopia japonica* by the coleopteran *Otiorhynchus sulcatus* (Curculionidae) during a greenhouse experiment.

The black vine weevil (*Otiorhynchus sulcatus*, Fig.1) is an exclusively parthenogenic coleopteran, meaning that adult weevils represent a single female clone (Smith, 1932; Son and Lewis 2005). This insect is widely dispersed all over the temperate regions and commonly encountered in Europe (Moorhouse et al. 1992). It is considered as a major pest of horticultural crops (Moorhouse et al. 1992) and Smith (1932) described more than 75 plant species, with observed herbivory in the USA. This high diversity of plant fed may lead to suppose that *O. sulcatus* could be a great predator for *F. japonica*.

**Fig.1.**
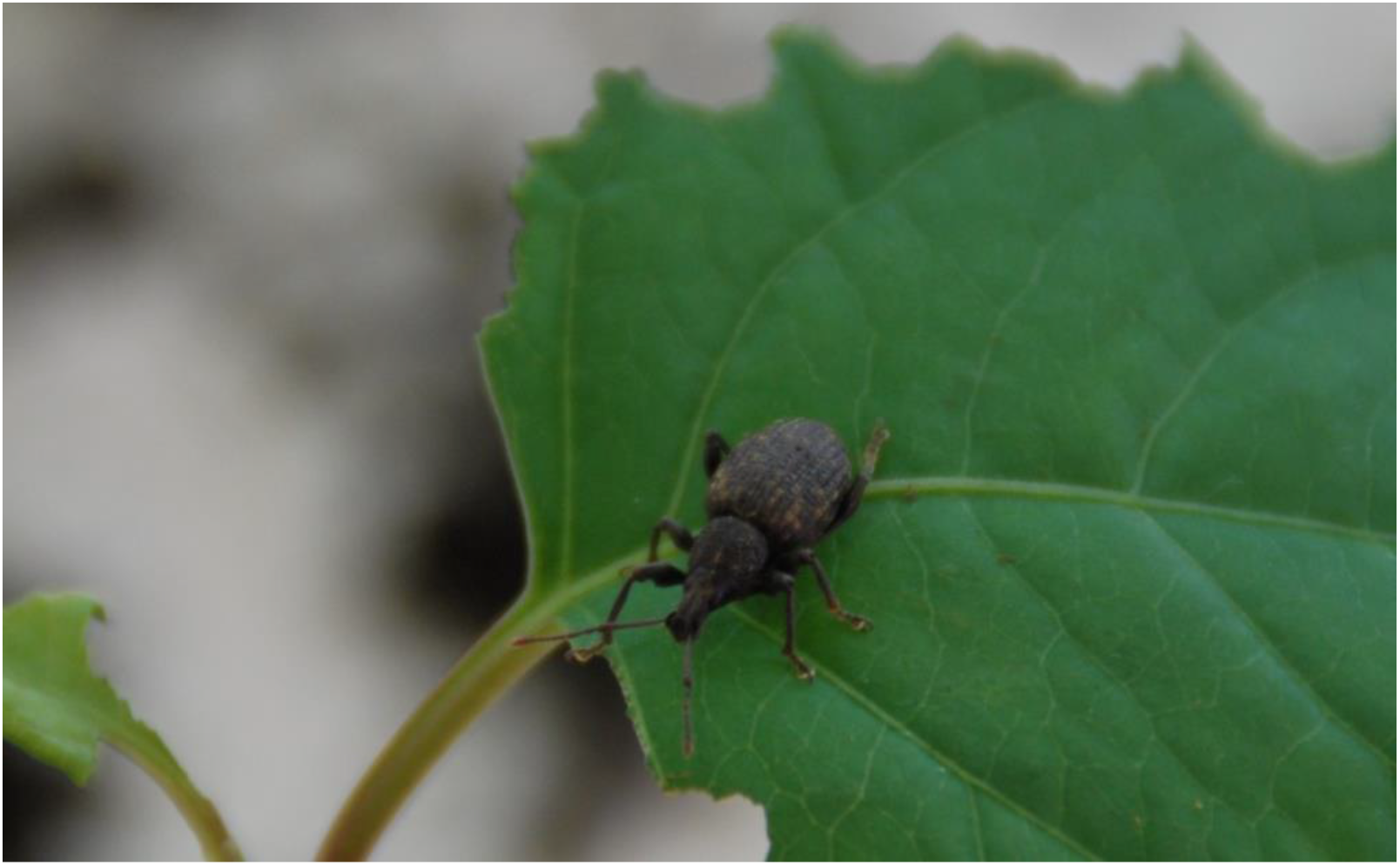
Picture of a Black vine weevil eating a Fallopia leaf. (© L. TEULIER)

To our knowledge, no direct field observation of herbivory of *O. sulcatus* on *F. japonica* were detected in France. And the lack of field description may be due to different parameters. *O. sulcatus* are indeed nocturnal, and field observation are obviously carried during the day. Another explanation could come from the low level of *O. sulcatus* population, because of the number of natural enemies (Moorhouse et al. 1992).

However, through an experimental protocol using *Fallopia* spp., herbivory of *O. sulcatus* was incidentally discovered. Japanese knotweed leaves were eaten, whereas other plants species present in the greenhouse stayed intact (personal observations). In order to characterize the origin of this potential herbivory, we found *O. sulcatus* larvae and some adults in each pot of *Fallopia*. We designed an experimental protocol to strengthen and validate this fortuitous finding. The goal of this study was therefore to characterize for the first time, the potential herbivory of *F. japonica* by a native coleopteran, *O. sulcatus* through two main questions:

a. Are *F. japonica* really eaten by *O. sulcatus*? Our hypothesis is that even if the insects were fed exclusively with *Fragaria*, they will prefer *F. japonica* when facing a 2-species choice. # Experiment 1 (“One night, one choice”)
b. If our latest hypothesis seems supported, are the insects only taste *F. japonica* because it is new and they met the opportunity? Our hypothesis is that *O. sulcatus* will eat the Japanese knotweed until the end. # Experiment 2 (“Until the End”)

## Material and methods

Insects, plants and assays were in the same conditions of climate room, at 23°C, 45-65% RH and a 12hr photoperiod / 24h.

### Insects

Two populations of *Otiorhynchus sulcatus* larvae were collected close to Lyon area in winter. The first population was collected in “la Doua” campus (Villeurbanne, France) in *Fallopia* pots, which is considered as the “F-Pop”. The second population of *Otiorhynchus* was collected in a horticultural nursery (Vienne, France) without *Fallopia* spp. It therefore constitutes the naive control population, called “N-Pop”. Both of these populations were reared to adults in climate room at 23°C, 45-65% RH and a 12hr photoperiod / 24hr. Weevil larvae were maintained on Strawberry plants (*Fragaria* spp.) in 1.3 L pots shut in vivarium or *F. japonica* until metamorphosis in adult (see Fisher and Bruck 2004).

### Plants

Rhizomes of *F. japonica* were collected from a single stand in Loire and stored at 4°C. Rhizome fragments were cut and selected with one node and a biomass of 1.5± 0.1 g. They were planted in FAVORIT^®^ peat soil and stored in a climate room (photoperiod = 16 hr light, 8 hr dark; temperature = 24°C) at Lyon University) until the plants have 7/8 expanded leaves for the 1^st^ assay and 5 expanded leaves for the 2d assay.

*Fragaria* spp. came from commercial plants.

### Experimental protocol

We decided to study only insect adult stage for focusing on the dramatic consequences on aerial herbivory in plants.

1. <u>“One night, one choice”</u>: insects from N-Pop (n=18) and F-pop (n=5) were randomly and individually placed in a Petri dish (diameter: 14.5 cm) filled with moist potting soil, in presence of one leaf piece of two plant species (*F. japonica* or *Fragaria*), following an experimental protocol adapted from Van Tol et al. 2004. Each leaf was previously cut into small rectangles of same size and scanned to obtain the initial area. 24h later, leaf area was measured and the difference between initial area and final area corresponded to the area eaten by the insects. As a control, for each day of experiment, a Petri dish with the two pieces of leaf without insect was placed in the same conditions.
2. <u>“Until the end”:</u> 15 adult black vine weevils from N-pop were randomly split into 5 2L-glass jars closed with plastic mesh, in presence of a plant of *F. japonica* (5 leafs each). Each day, pictures were taken to constitute a time-lapsed evolution of herbivory. The experiment ended when all the leaves were eaten.

## Results and Discussion

### Fallopia is attractive, even for naive herbivores

*Fallopia*-rearing (F-pop) but also naive (N-pop) insects were able to eat both of the plant species (Fig. 2, Table 1). Indeed, insects have eaten a significant leaf area during the night of experiment (between 0.99 and 1.70 cm^2^/night). F-pop *O*.*sulcatus* ate a higher percentage of *Fallopia* than *Fragaria* (80% vs. 20%, respectively). N-pop weevils behaviour was dependent on what they ate before the experiment. Their choice seems to be driven by the novelty and therefore the attractiveness between *Fragaria* and *Fallopia* appears more balanced than for F-pop insects. N-pop insects reared on *Fragaria* were also more attracted by *Fallopia* than *Fragaria*, and on the opposite, N-pop weevils reared on *Fallopia* ate more *Fragaria* than *Fallopia* leaves (67.7% vs. 32.3%). Moreover, on the whole sampling size of 41 tests, 10 weevils ate *Fallopia* only, whereas only 2 ate *Fragaria* only. The 29 others tried both of the plants, which supposed that for a major part of them, it is not a randomized choice.

**Fig.2.**
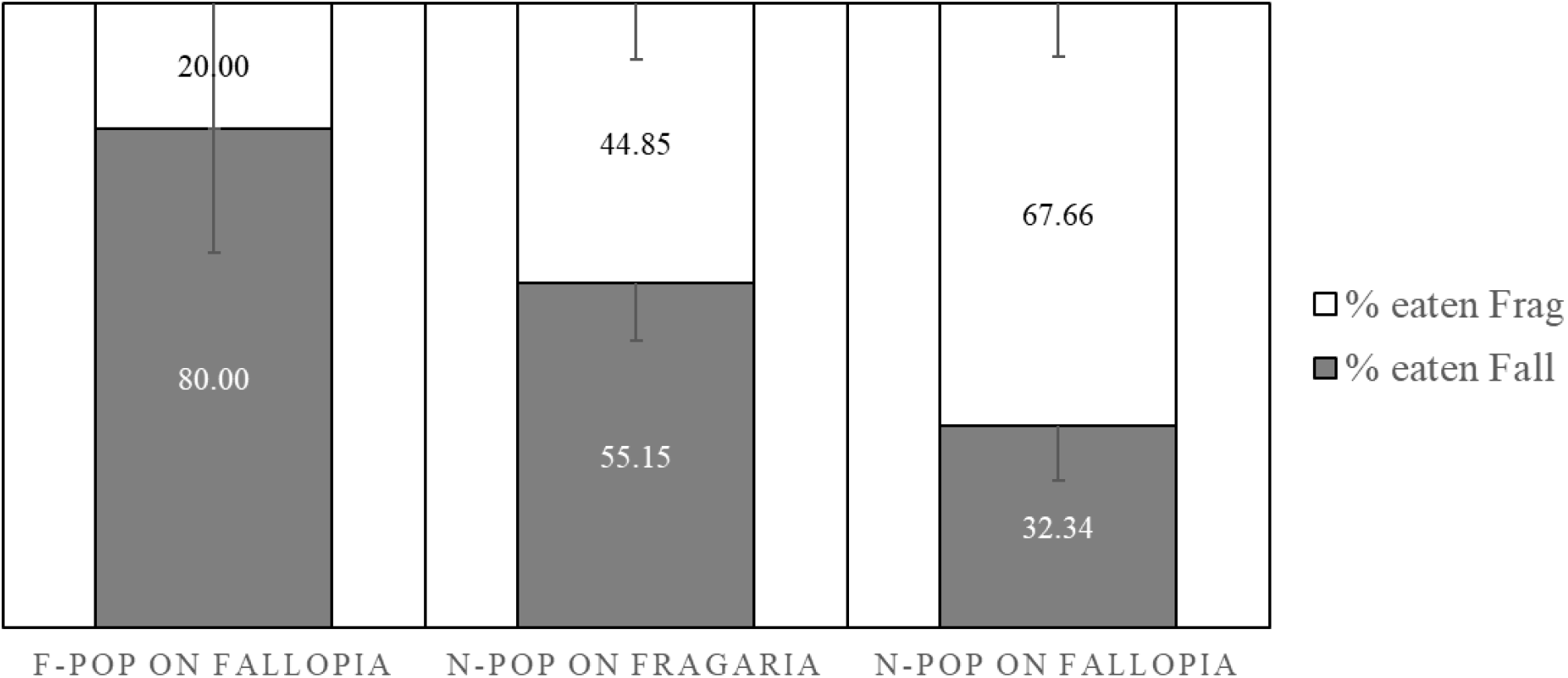
Percentage of *Fallopia* (grey) or *Fragaria* (white) leaf area eaten by black vine weevils during one night. Each insect was able to choose between the same leaf area (3-10 cm^2^) of Japanese knotweed and strawberry. F-pop on *Fallopia* (n=5) means for the weevils coming from the experimental greenhouse, found in *Fallopia* pots, N-pop on *Fragaria* (n=18) and N-pop on *Fallopia* correspond to naïve weevils sampled in the commercial greenhouse, without Japanese knotweed and reared on *Fallopia* or on *Fragaria*, respectively. For more details, please refer to Materials and Methods section.

**Table 1.**
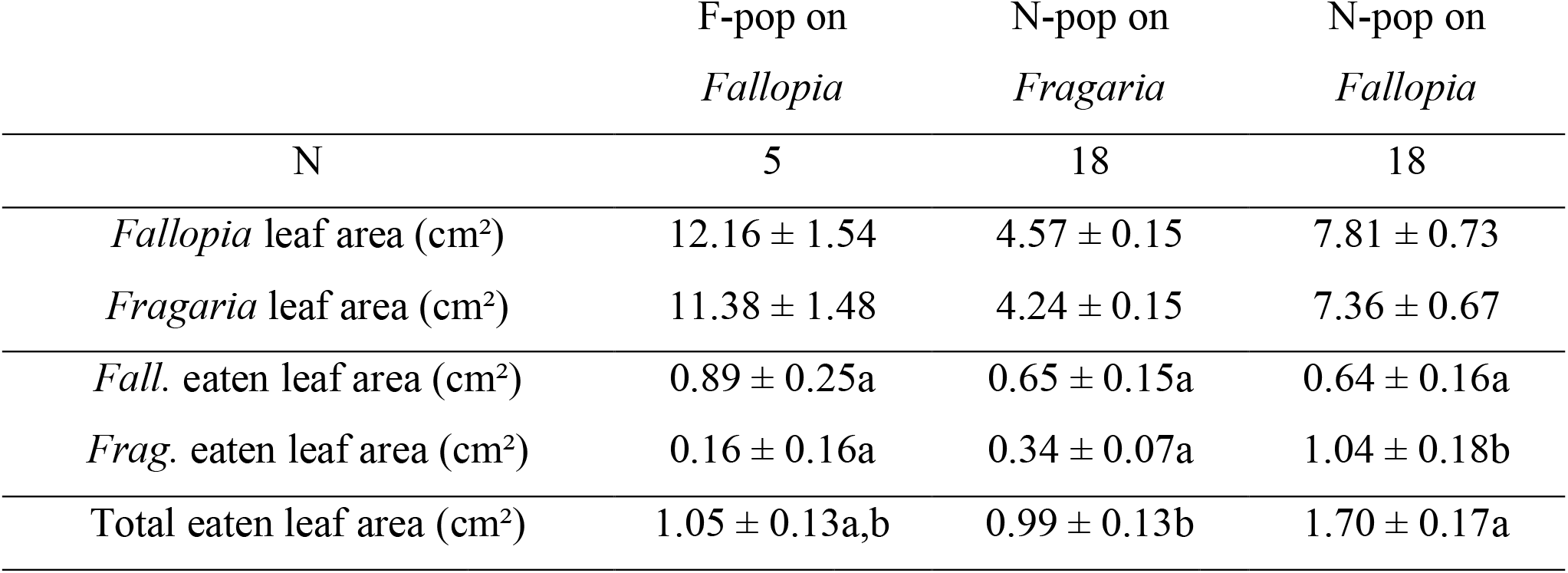
Characteristics of leaves used for the experiment ‘One night, one choice’. Insects found in the experimental greenhouse of the Campus La Doua (Villeurbanne, France) are named ‘F-pop’, whereas the ‘N-pop’ represents the naïve insects collected close to Vienne (∼30km of Lyon, without any *Fallopia* around), reared on *Fragaria* or *Fallopia*.

### Not only a test

One may argue that Black wine weevils, which preferred *Fallopia* in the former experiment, only tasted the Japanese knotweed, but would not like it. This hypothesis seems unlikely, because of the presence of weevil larvae and the obvious herbivory of adults exclusively in the *Fallopia’s* pots, even if there was a lot of other plants available in our experimental greenhouse. To confirm the hypothesis that *Fallopia* spp. are sufficiently appetent, Black wine weevils ate the whole plant in ∼25days, until no leaf left (Fig. 3).

**Fig.3.**
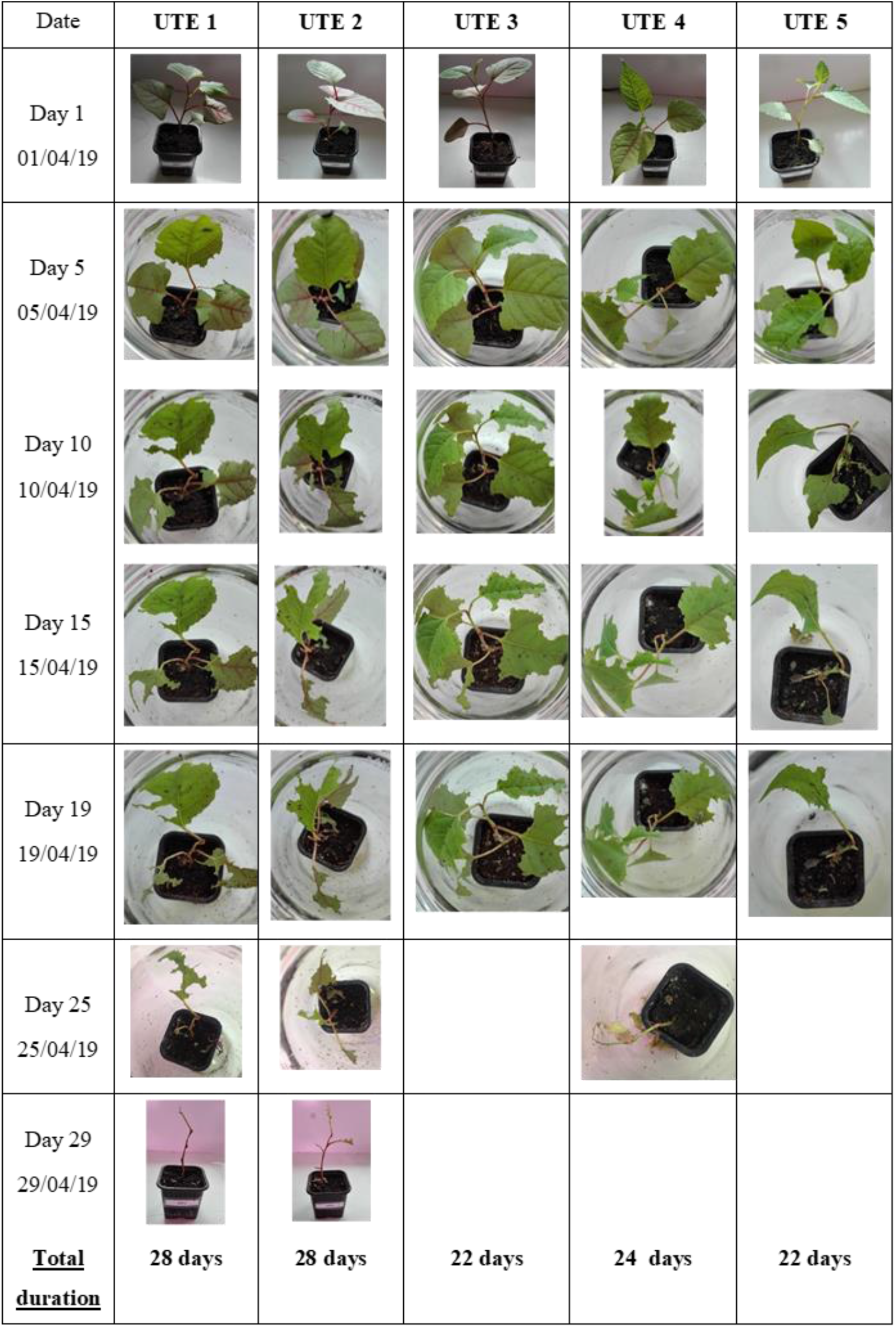
Series of pictures showing the evolution of 5 *Fallopia* pots in presence of 3 black wine weevil adults. They ate all the leaves in ∼25 days.

## Conclusion and perspectives

For the first time, our results clearly show a massive herbivory of the invasive Japanese knotweed in France by an insect: *Otiorhynchus sulcatus*. According to the enemy release hypothesis (Keane and Crawley 2002), introduced species can experience less selective pressures from natural enemies. Indeed, *Fallopia* has very little herbivory (Ness et al. 2013; Gippet et al. 2018) while in its native zone, *F. japonica* is heavily impacted by herbivores (Kawano et al. 1999). Nevertheless, it is possible to envisage an evolution in the territory of invasion leading to the adaptation of native herbivores to the *Fallopia* taxon.

The black vine weevil, *O. sulcatus* is a polyphagous insect that is a nocuous pest of field and is cited as one of the most important species afflicting crops globally throughout the United States, western Canada, and northern Europe (Moorhouse et al. 1992). Even if this simple observation may appear as trivial, it highlights a novel plant-insect interaction and may start new insight in plant control or invasion management. Indeed, depending on the point of view, this interaction could be considered as a serious asset for biological control of the Japanese knotweed through the herbivory of a native insect, contrary to other insects specifically introduced, such as *Gallerucida bifasciata* (Wang et al., 2010a) or *Euops chinesis* (Wang et al. 2010b), which failed to be efficient only on this target. It could be also considered as a starting point of a pest control method, using *Fallopia* as a trap plant in greenhouse to catch black wine weevils. These latter points need further investigations for their validation.

## Declarations

This study was supported by the recurrent funding of the Claude Bernard University Lyon1 attributed to LT and FP. The authors declare no conflict of interest. LT and FP designed the experimental protocol. SP and CB performed the experiments. All of the authors participated to draft and review the manuscript.

## Acknowledgments

We are grateful to the reviewers for their further comments on our manuscript. We would like to thank Tatiana Buisson and Rémi Bernard for their help during the experiments. We thank Franck Poly (Sempervivum et Cie nursery) for providing us *O. sulcatus* and Bernard Kaufmann for the insect identification.

## Notes

### Competing Interest Statement

The authors have declared no competing interest.

